# SpotSweeper: spatially-aware quality control for spatial transcriptomics

**DOI:** 10.1101/2024.06.06.597765

**Authors:** Michael Totty, Stephanie C. Hicks, Boyi Guo

**Author notes:** Corresponding author: Boyi Guo.

## Abstract

Quality control (QC) is a crucial step to ensure the reliability and accuracy of the data obtained from RNA sequencing experiments, including spatially-resolved transcriptomics (SRT). Existing QC approaches for SRT that have been adopted from single-nucleus RNA sequencing (snRNA-seq) methods are confounded by spatial biology and are inappropriate for SRT data. In addition, no methods currently exist for identifying histological tissue artifacts unique to SRT. Here, we introduce SpotSweeper, spatially-aware QC methods for identifying local outliers and regional artifacts in SRT. SpotSweeper evaluates the quality of individual spots relative to their local neighborhood, thus minimizing bias due to biological heterogeneity, and uses multiscale methods to detect regional artifacts. Using SpotSweeper on publicly available data, we identified a consistent set of Visium barcodes/spots as systematically low quality and demonstrate that SpotSweeper accurately identifies two distinct types of regional artifacts, resulting in improved downstream clustering and marker gene detection for spatial domains.

## 1 Main

Spatially-resolved transcriptomics (SRT) has revolutionized our ability to profile cells in their spatial context, providing unprecedented insights into human health and disease. This technology not only enables the exploration of cellular heterogeneity and interactions within defined tissue architectures, but it also catalyzes the advancement of computational tools designed for SRT data analysis [1–3]. Many computational tools have been designed to improve or augment existing workflows designed for single-cell analysis, including spatially variable gene detection [4], spot-level cellular deconvolution [5], and spatially-aware clustering [6]. While the transition from single-cell data analysis to spatially-aware computational strategies has enhanced the resolution of biological inference by using spatial information, one critical aspect, namely quality control (QC), has been overlooked.

For next generation sequencing technologies, QC is a process that helps identify and remove low quality observations which may negatively impact downstream analyses, such as clustering and differential expression tests, leading to spurious findings [7, 8]. Unlike single-cell/nucleus RNA-sequencing (sc/snRNA-seq), which capture mRNA transcripts from a cell body or nucleus, SRT profiles mRNA transcripts from a wide variety of biological domains (i.e., neuronal processes vs cell bodies) that display substantial variation in gene expression signatures [8]. However, current methods for detecting outliers or low quality observations in SRT use methods developed for sc/snRNA-seq, such as fixed and data-driven global thresholds, which implicitly assume that all observations are derived from a homogeneous sample (i.e., exclusively cell bodies). We show here that these methods fail to account for biological heterogeneity present in SRT and result in unwanted biases at the stage of QC. For example, in human brain tissue, global QC methods naively flag more low quality observations from white matter compared to gray matter due to the natural molecular and cellular differences [8–11]. As spatial atlases increasingly grow in size [12], this motivates the need to develop robust, spatially-aware QC methods to ensure the integrity of downstream analyses using SRT data.

Here, we introduce spatially-aware QC metrics and a computational pipeline to identify and discard low-quality observations and regional artifacts generated by sample processing errors in SRT data. We illustrate the utility of our methods on postmortem human brain tissue with expert manual annotations profiled on the 10x Genomics Visium Spatial Gene Expression platform [9, 10]. We first demonstrate that standard QC metrics are confounded with natural biological heterogeneity. Compared to widely used global QC methods, our spatially-aware QC approach is less susceptible to these biological confounds which enables the preservation of high-quality spots across diverse spatial domains, thus ensuring the integrity of downstream analyses. Applying SpotSweeper to multiple publicly available datasets, we identified a set of Visium barcodes that display systematically low library size. Moreover, using multiscale approaches, we demonstrate that SpotSweeper is able to accurately identify two distinct classes of regional artifacts within the tissue, namely dryspots and hangnails, caused by incomplete coverage of permeabilization agents and tissue damage, respectively. These methods are implemented in the SpotSweeper R package within the Bioconductor framework, allowing for direct integration with workflows using established Bioconductor infrastructure for SRT data [13].

## 2 Results

### 2.1 Overview of SpotSweeper and the methodological framework

The SpotSweeper framework introduces two spatially-aware QC approaches for SRT data that can identify (i) individual low quality spots and (ii) region-level artifacts in a tissue section across multiple spots. We utilize established QC metrics such as library size or total unique molecular identifiers (UMI) [14], number of unique genes detected [15], and percent of reads mapping to mitochondrial genes [16] for both spot-level and regional artifact detection.

We first introduce the spot-level QC approach (**Figure 1A**) based on established methods for spatial outlier detection [17]. For each spot *i*, we define a local neighborhood using *k*-nearest neighbors based on spatial coordinates around each spot. Then, we calculate a robust *z*-score for all spots in the neighborhood:

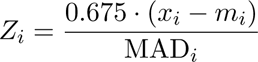

**Figure 1:**
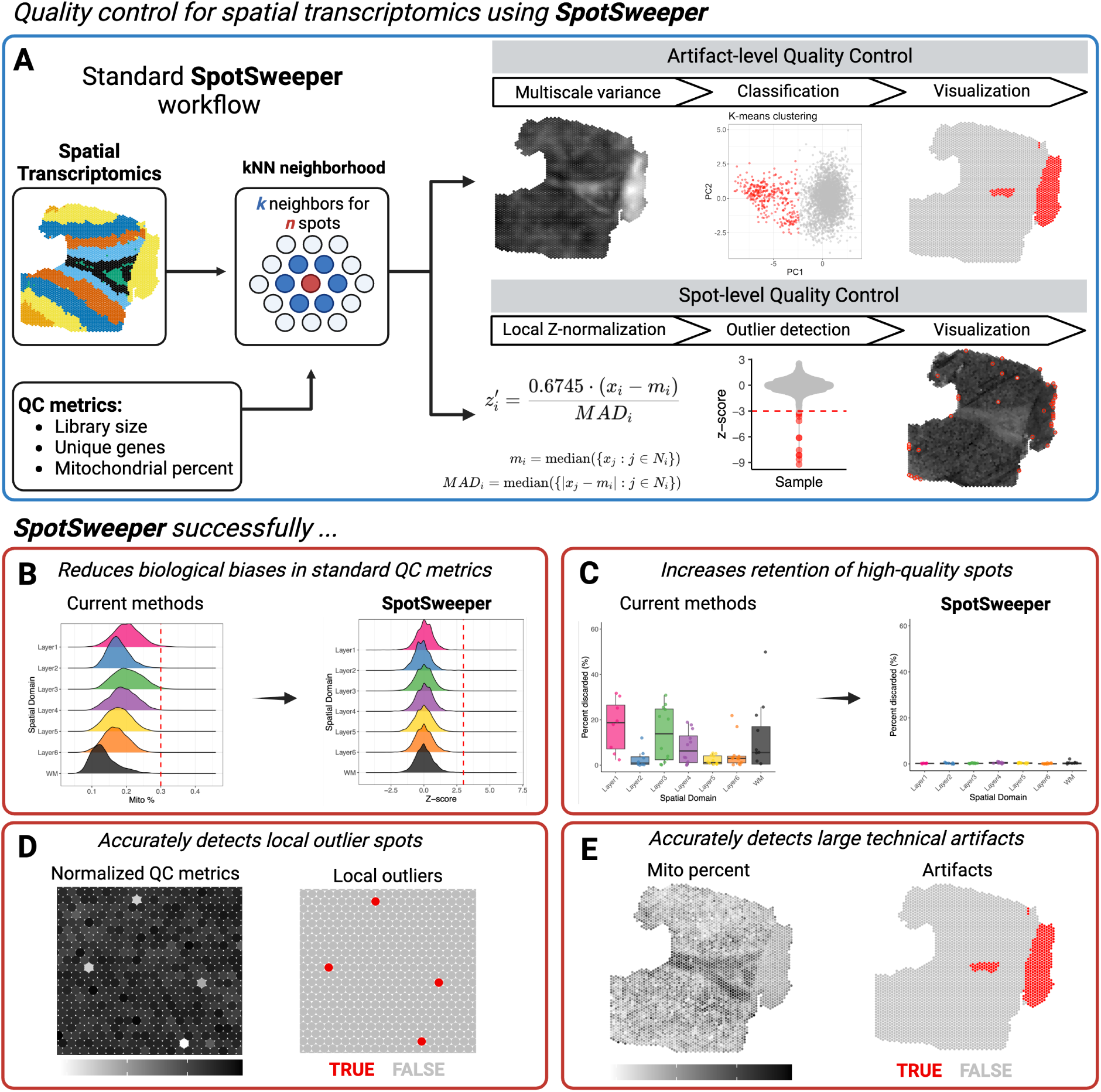
An overview of SpotSweeper: spatially-aware QC methods to identify and eliminate low-quality spots and region-level artifacts in SRT data. Data: postmortem human brain tissue section profiled on the 10x Genomics Visium Spatial Gene Expression platform with annotated spatial domains for gray matter (Layers 1-6) and white matter (WM) [10]. (A) Using the *k*-nearest neighbors of each spot, SpotSweeper identifies region-level artifacts and low-quality spots. In contrast to exisiting QC metrics for SRT data, key advantages of SpotSweeper are it (B) is less biased by differences across spatial domains, (C) retains more high-quality spots, (D) accurately detects local outliers, and (E) accurately detects compromised spots due to region-level artifacts. Created with BioRender.com

where *x_i_*is the QC metric (e.g. library size, number of detected genes, or the percent of reads mapping to mitochondrial genes) for the *i*th spot, *m_i_* is the median of the neighbors’ values, and the denominator is the median absolute deviation (*MAD_i_*), defined as:

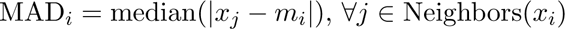

We add a scaling factor of 0.675 (the 75th percentile of the standard normal distribution) to make the MAD comparable to the standard deviation under the assumption of normally distributed data [18]. This in turn makes the proposed *z*-score comparable to a standard *z*-score. Spots can then be discarded as local outliers based on their *z*-score.

Next, we introduce the region-level QC approach (**Figure 1A**). Our method is based in the idea that region-level artifacts can be distinguished by unusually small variation in mitochondrial ratio due to loss of natural biology variability and we give examples in the following sections. To enhance the detection of these artifacts, we implement a multiscale approach that leverages multiple scales or varying neighborhood sizes to capture both local and broader spatial patterns. Similar to the spot-level QC methods, the *k* - nearest neighbors for each spot are first identified based on the spatial coordinates. For each neighborhood size (i.e., scale), local variance of the mitochondrial ratio is calculated for each central spot and adjusted for a mean-variance relationship using robust linear regression via the iterative re-weighted least squares algorithm (see **Methods** for more details). The residuals of the linear regression are taken to be the mean-corrected local variance. Principal component analysis is then performed on the mean-corrected local variances of all neighborhood sizes for dimensionality reduction. *k*-means clustering (*k*=2) is then performed in the first two principal components to identify regional artifacts compared to high-quality tissue.

### 2.2 Key innovations of SpotSweeper

The key innovations of SpotSweeper compared to commonly used QC approaches for SRT data are as follows. First, SpotSweeper assesses the quality of individual spots relative to their local neighborhood as opposed to existing approaches that assess the quality of spots relative to spots across the whole tissue slide. This is implemented using *k*-nearest neighbors via spatial coordinates and helps overcome potentially problematic global QC approaches due to differences in spatial domains (**Figure 1B**) and retains more high quality spots (**Figure 1C**). Second, SpotSweeper leverages local outlier approaches leading to improved spatially-aware QC metrics within a tissue (**Figure 1D**), and can be applied across multiple tissue sections. Finally, SpotSweeper is the first method capable detecting of region-level artifacts that are distinct in SRT data by taking advantage of the local variance of QC metrics (**Figure 1E**).

### 2.3 Global QC approaches are confounded with spatial domains

In this section, we show that standard QC metrics are confounded by natural biological variation in SRT. Consequently, commonly used QC approaches that identify global outliers across entire tissue sections lead to biased removal of spots across spatial domains. Here, we use a dataset profiling dorsolateral prefrontal cortex (DLPFC) from postmortem human brain tissue measured on the 10x Genomics Visium Spatial Gene Expression platform [10]. We picked this dataset because the DLPFC contains substantial differences across spatial domains, namely between white matter (WM) and six gray matter domains (cortical layers L1-L6) (**Figure 2A**). Some layers, such as L2, L3, L5, and L6 contain cell-bodies (i.e., soma), while other layers (L1 and WM) exclusively contain neuronal processes (i.e., dendrites and axons). In addition, soma-rich layers substantially differ in cell-type composition.

**Figure 2:**
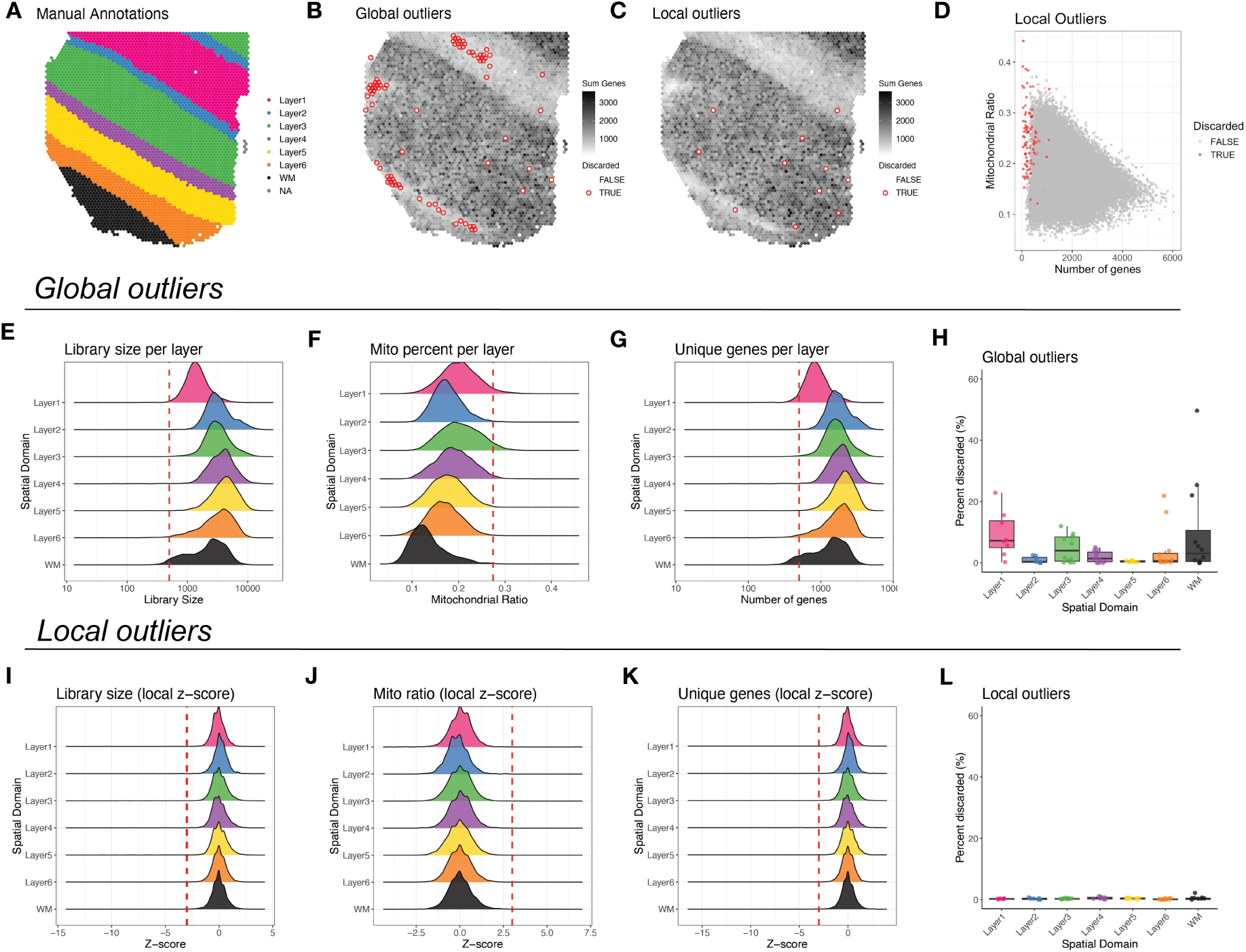
SpotSweeper improves quality control using local versus global approaches to identify outliers or low quality spots. (**A**) Manually-annotated cortical layers in a single human dorsolateral prefrontal cortex (DLPFC) tissue sample from Maynard et al. [10]. (**B-C**) Spot plots displaying the number of detected genes overlaid with the low quality spots (red) identified using (**B**) common QC approaches across the tissue (global outliers) using fixed thresholds (*>*0.275 and *<*500, respectively) and (**C**) local outliers as detected by SpotSweeper. (**D**) Scatter plot of the number of detected genes (*x*- axis) and percent of reads mapping to mitochondrial genes (*y*-axis) with low quality spots identified using SpotSweeper. (**E-G**) Ridge plots of the distribution of library size, percent of mitochondrial genes, and number of detected genes across cortical layers. Red dotted lines indicate fixed thresholds to identify outliers. (**H**) Box plots of the percent of discarded spots (global outliers) across cortical layers (*n*=12 tissue samples). (**I-K**) Ridge plots showing the distribution of *z*-normalized QC metrics for library size, mitochondrial ratio, and unique genes. (**L**) Box lots displaying the percent of discarded spots (local outliers) across cortical layers using SpotSweeper.

Current approaches typically perform spot-level QC based on approaches developed for snRNA-seq data [19, 20], namely, setting global fixed [15] and data-derived thresholds [21–23]. Using standard QC metrics for SRT data, such as library size, proportion mitochondrial genes, and number of unique genes [19, 20] we show here that these global QC approaches result in an uneven number of spots being labeled as low quality across spatial domains. (**Figures 2B**). As expected, soma-rich layers (L2-6) showed greater library sizes and number of unique genes compared to spatial domains that only contain neuronal processes (L1 and WM) (**Figures 2E,G**), whereas L1 and L3 showed the highest mitochondrial ratio (**Figure 2F**). In fact, global outliers detected with moderately conservative fixed thresholds set at 500 total genes, 500 unique genes, and 0.275 mitochondrial percent were biased to discarding spots in layers with low number of transcripts (L1 and WM) and high mitochondrial ratio (L3) when applying SpotSweeper to multiple tissue sections within the Maynard et al. [10] DLPFC samples (*n*=12 tissues) (**Figures 2H, S1**). This resulted in an average of 9.34%, 4.70%, and 9.74% of spots excluded from L1, L3, and WM, respectively, across samples.

### 2.4 QC approaches based on local outlier detection controls for confounding biology

Using SpotSweeper, we show that by restricting outlier detection to local neighborhoods, our approach reduces the biased exclusion of spots across different spatial domains (**Figure 2C**), while still identifying spots with relatively low library size/unique genes and high mitochondrial ratio (**Figure 2D**). We show that *z*-normalizing QC metrics based on local neighborhoods successfully normalizes their distributions across spatial domains (**Figures 2I-K**). For defining local outliers, we chose to use a cutoff of three standard deviations from the mean under a standard Normal distribution. Using thresholds of *<*-3 *z*- scores for library size and unique genes detection, and *>*3 *z*-scores for mitochondrial ratio, this approach leads to discarded spots more uniformly distributed across the spatial domains compared to global QC approaches using either using fixed thresholds (**Figure 2L**), or data-driven thresholds such as median-absolute deviations (**Figure S1**). SpotSweeper excluded an average of 0.21%, 0.28%, and 0.40% of spots excluded from L1, L3, and WM, respectively, which ultimately resulted in the retention 1,670 high quality spots (an average of 139.17 per sample) compared to fixed thresholds.

### 2.5 SpotSweeper detects consistent set of spots with systematically low library size

When applying SpotSweeper to the Maynard et al. [10] DLPFC samples (*n*=12 Visium samples), we noticed SpotSweeper identified a consistent set of six spots as low quality based on library size across all 12 tissue sections (**Figure 3A**). This motivated us to expand the datasets considered to explore if additional datasets also identified a similar set of low quality spots. We considered a larger DLPFC data from Huuki-Myers et al. [9] with *n*=30 Visium samples as well as *n*=1 Visium samples of mouse coronal brain sections generated by 10x Genomics. In all three datasets, we found that the identical six spots contained less total UMI counts (or library size) compared to neighboring spots (indicated by negative *z*-scores) in all samples across all three datasets (*n*=43 total) (**Figure 3B**), and were considered local outliers (*<*-3 library size local *z*-scores) in over half of all samples (**Figure 3C**). The total UMI counts, unique genes, and mitochondrial ratio for these spots versus all others are shown in **Figure S2**. Only spots underlying tissue samples were included in these analyses. Importantly, we did not find any spots that with higher than average transcripts compared to neighbors that were repeatedly detected as local outliers across many samples (**Figure 3C**).

**Figure 3:**
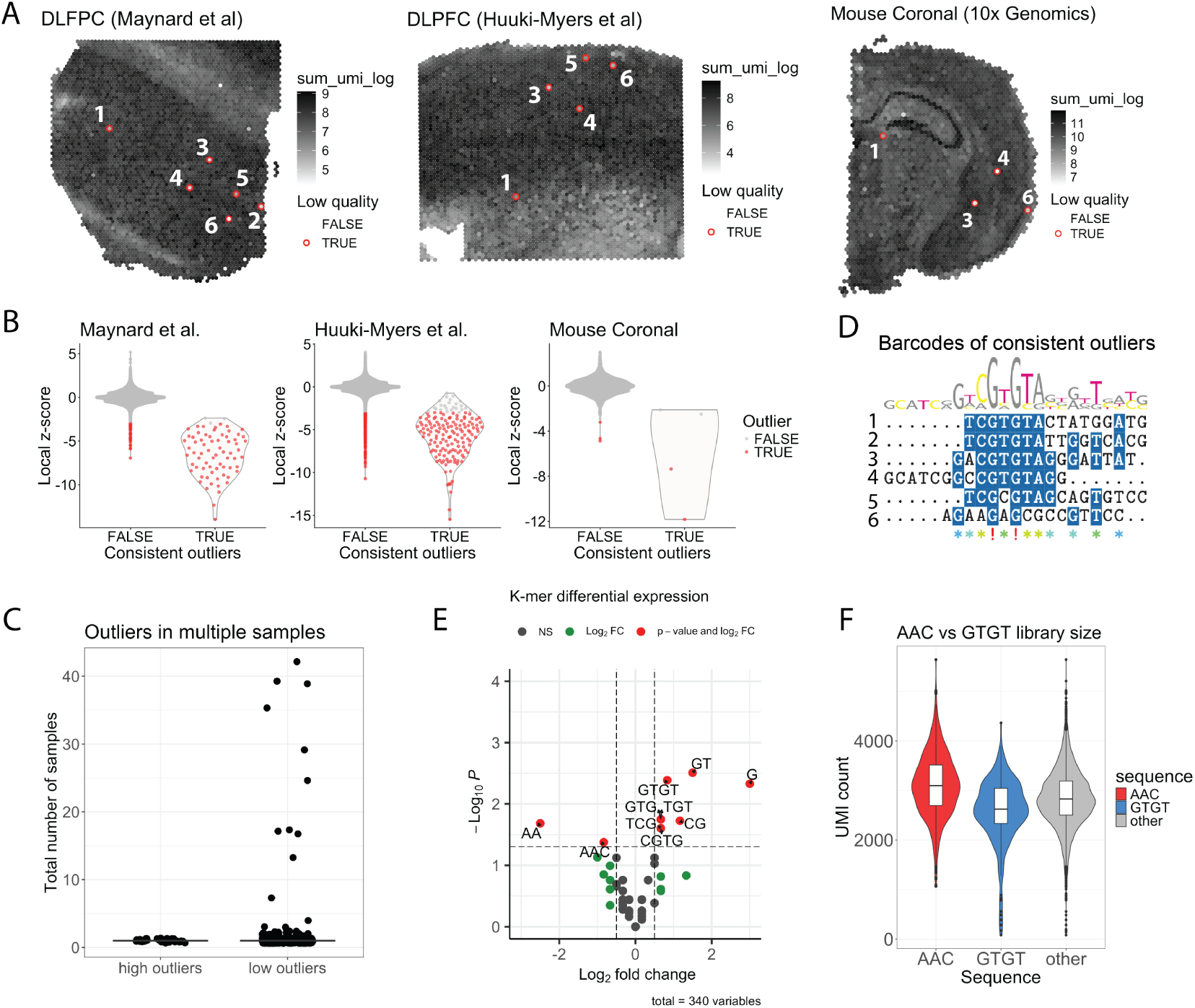
SpotSweeper detects consistent set of spots with systematically low library size driven by barcode biases. (**A**) Six Visium barcodes/spots were consistently flagged as having systematically-low library size Maynard et al. [10] (*n*=12), Huuki-Myers et al. [9] (*n*=30), and mouse brain (10x Genomics; *n*=1) datasets. (**B**) Violin plots comparing local library size *z*-scores for consistent outliers versus all other spots for each dataset. Red dots were detected as outliers by SpotSweeper. (**C**) Spots detected as local outliers based on high library size (*>*3 *z*-scores) are not found across multiple samples, unlike low outliers (*<*-3 *z*-scores). (**D**) Best-fit sequence alignment of the DNA barcodes underlying consistent outliers shows substantial homology with 4 out of 6 barcodes containing a CGTGTA sequence. (**E**) Volcano plot showing differentially expressed *k*-mer sequences between consistent outliers and the top six barcodes ranked by mean library size across all Visium samples (*n*=43). Positive values indicate increased expressing in outlier spots. (**F**) Boxplots of total UMI counts for Visium barcodes/spots that contain differentially expressed *k*-mers from top ranked (AAC) and outlier (GTGT) spots show biases towards higher and lower library sizes, respectively, compared to all other spots.

In the Visium platform, every spot has a synthetic DNA barcode that is assigned to a specific spatial coordinate, and the barcode assigned to a given spatial coordinate is same across all Visium assays. Considering previous work demonstrating how synthetic DNA barcodes sequences can lead to downstream bias in PCR amplification [24, 25], we hypothesized that the synthetic DNA barcodes associated with these problematic Visium spots are responsible for downstream biases. If so, we predicted that barcodes likely have homologous sequences. To this hypothesis, we performed multiple sequence alignment of these six barcodes and indeed found remarkable homology with four out of six barcodes containing a CGTGTA sequence (**Figure 3D**). To determine if barcode sequences may be driving downstream biases in library size, we next conducted differential *k*-mer analysis between these six low quality spots and the top six mean-ranked barcodes (**Figure S3**) to determine if there were *k*-mers differentially associated with barcodes that consistently show small or large library size, respectively. We found that a number of *k*-mers were indeed differentially expressed between the six consistent outliers compared to the six top mean-ranked spots (**Figure 3E**), and these differentially expressed *k*-mers were sufficient to distinguish outlier from top mean-ranked barcodes (**Figure S3**). AAC and GTGT were the longest *k*-mers associated with top ranked and low quality spots, respectively. We finally show that, on average, spots containing AAC and GTGT *k*-mers showed bias towards larger and smaller library sizes, respectively (**Figure 3F**). Collectively, these results suggest that the spatial barcodes present in Visium arrays have inherent biases that lead to systematically low library size and unique genes detected.

### 2.6 Spatial transcriptomics methods are susceptible to regional artifacts

In histopathology, tissue artifacts have been defined as “an artificial structure or tissue alteration on a prepared microscopic slide as a result of an extraneous factor” [26] and have been characterized to include (pre)fixation, processing, staining, or mounting artifacts [27]. In the context of the 10x Genomics Visium Spatial Gene Expression platform, tissues are also mounted on slides and regional artifacts can occur in a similar way. However, there currently lack studies both characterizing and identify regional artifacts. In this section, we begin by considering the Huuki-Myers et al. [9] dataset with *n*=30 Visium samples and characterize two types of regional artifacts unique to SRT data (**Figure 4**). Finally, we demonstrate how SpotSweeper provides computational methods to detect the tissue artifacts (**Figure 5**).

**Figure 4:**
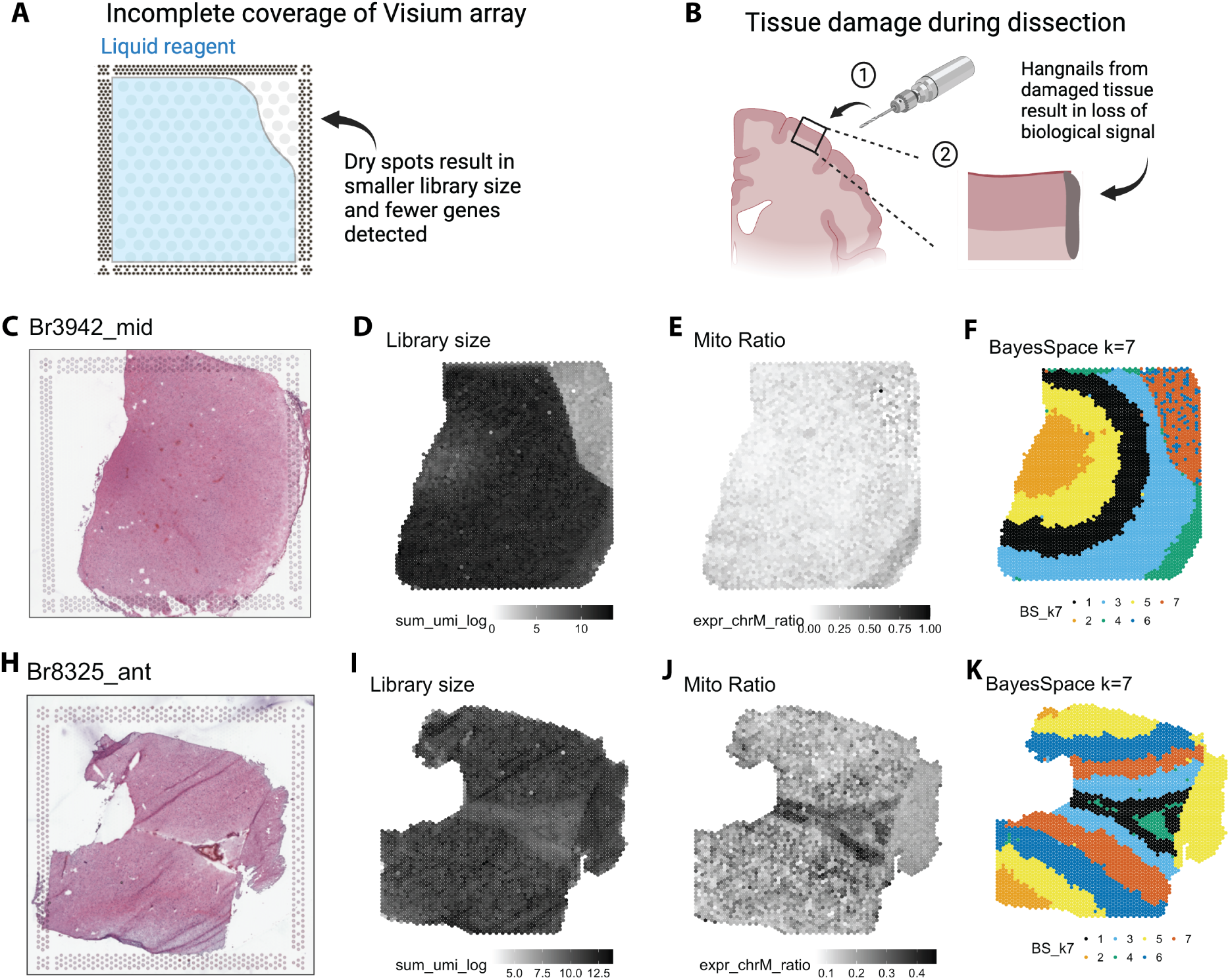
Characterizing region-level technical artifacts unique to spatial transcriptomics. Schematics showing how (**A**) incomplete coverage of Visium capture areas by permeabilization liquids (i.e., dry spots) results in large regions with low library size and unique genes, and (**B**) tissue damage during (1) dissection from a drill used to dissect human brain regions results in (2) hangnail artifacts with attenuated or altered biological signal. Dryspot artifacts (**C**) cannot be seen in histological images, but instead present as areas with (**D**) low library sizes and (**E**) no difference mitochondrial ratio. (**F**) Dryspot artifacts distinctly cluster using spatially-aware clustering methods. (**H-J**) Unlike dryspots, hangnail artifacts are clearly visible in histological images, present with similar library size and mitochondrial ratio as the rest of the sample, and get incorrectly clustered with one of the cortical layers. In this case, BayesSpace cluster 5 which we approximate as cortical layer 6. Panels A and B created with BioRender.com

**Figure 5:**
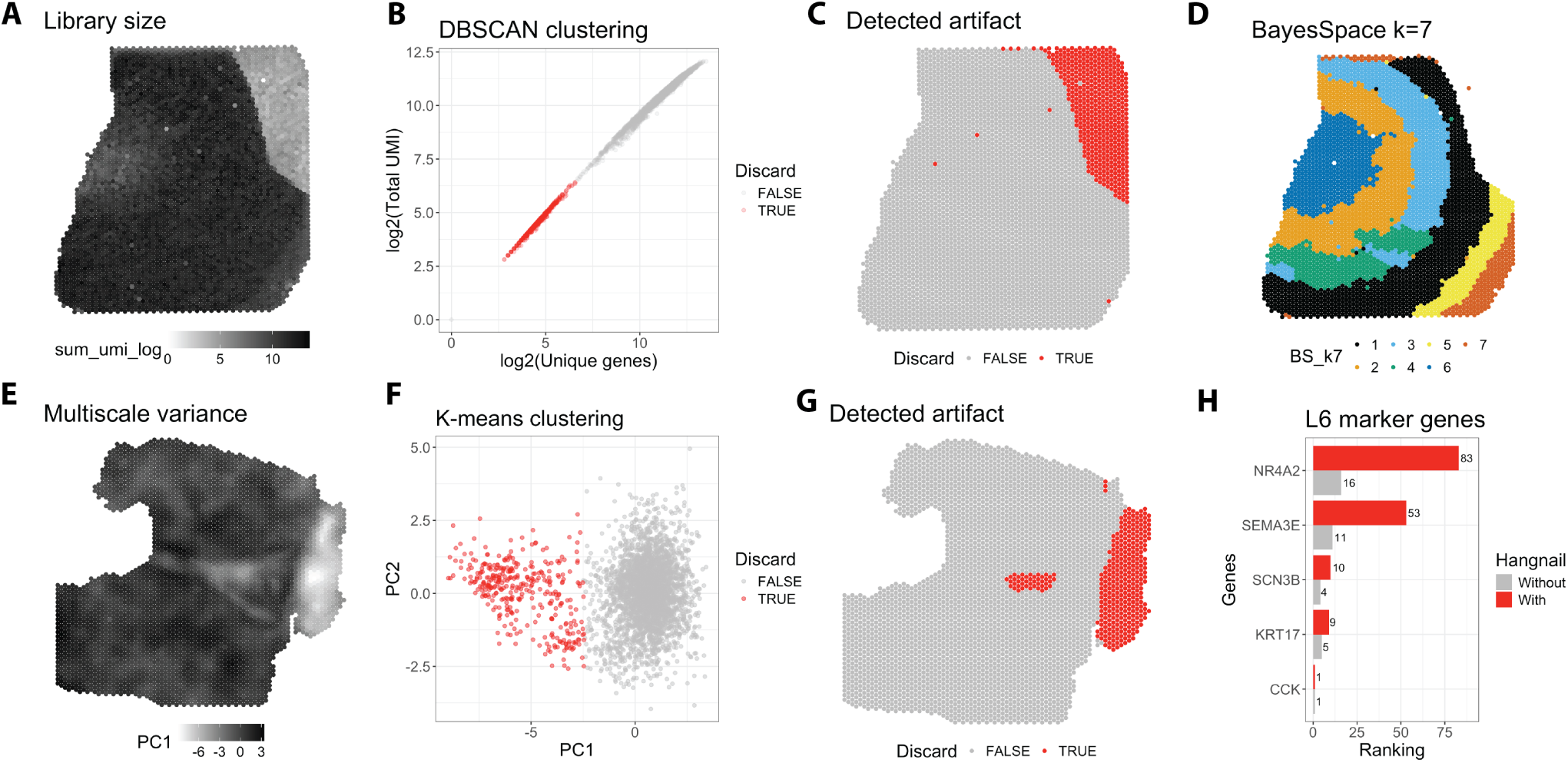
SpotSweeper accurately detects dryspot and hangnail tissue artifacts. (**A**) Spot plot visualizing a dryspot artifact using log2-transformed library size. (**B**) Density-based clustering methods (DBSCAN) can be used to detected dryspot tissue artifacts in log2-transformed library size and unique gene space. (**B**) Spot plots showing dryspot artifact automatically annotated as the cluster with smaller library size. (**D**) Removing dryspot artifacts results in the discovery of an additional spatial domain using spatially-aware clustering (BayesSpace). (**E**) Spot plot displaying the first principal component of the multiscale variance (1-5 concentric circles for each spot) of mitochondrial ratio. (**F**) *k*-means clustering (*k*=2) on the first two principal components of the multiscale variance successfully distinguishes the hangnail artifact from high-quality tissue. (**G**) Spot plots showing the hangnail artifact automatically annotated as the cluster with lower average multiscale variance. (**H**) Bar plots showing that removing the detected hangnail increases the ranking of canonical L6 marker genes.

The first type of region-level artifact is an incomplete coverage of permeabilization agents, and referred to here as ‘dryspot’ artifacts. These dryspot artifacts are caused by incomplete tissue coverage of permeabilization agents (**Figure 4A**). Permeabilization is an essential step that releases mRNA content from tissue samples that are then captured and barcoded in each Visium spot. Because dryspot artifacts are not due to differences in tissue quality (**Figure 4C**), they present as regions with drastically lower library size (**Figure 4D**) and unique genes (**Figure S4**), but no substantial difference in mitochondrial ratio (**Figure 4E**). Due to this drastic difference in detected transcripts, dryspots often form distinct clusters using spatially-aware clustering methods (**Figure 4F**). This prevents the accurate detection of six cortical layers and white matter in the DLPFC tissue section, and further confounds the underlying biology.

The second type of artifact is from tissue damage that may occur during dissection, and referred to here as ‘hangnail’ artifacts. These artifacts are caused by tissue damage, such as from a high-powered drill used to dissect small regions from a frozen human brain (**Figure 4**). This form of tissue damage results in loss of interpretable biological signal. Unlike dryspot artifacts, hangnails are often visually apparent in histological images (**Figure 4H**), but do not show substantial differences in average library size (**Figure 4I**), genes detected (**Supplementary Figure S4B**), or mitochondrial ratio (**Figure 4J**). Because of this, spots underlying hangnail artifacts tend to cluster with spatial domains (in this case, L6) corresponding to high-quality, non-damaged tissue regions (**Figure 4K**). This presents a significant problem for artifact removal.

### 2.7 SpotSweeper identifies regional artifacts unique to spatial transcriptomics

Here, we propose methods to identify region-level artifacts, both dryspot and hangnail artifacts. Because dryspot artifacts present with substantially lower library size (**Figure 5A**), Huuki-Myers et al. [9] previously discarded this dryspot using manual library size thresholds. However, we show in Figure 2 that fixed thresholding is biased by differences across biological domains and results in the inadvertent discarding of high quality tissue spots. SpotSweeper improves this by implementing DBSCAN clustering on log2-normalized QC metrics of library size and number of unique genes detected to identify dryspot artifacts (**Figures 5B-C**). Removing the dryspot artifact improves spatial domain detection, allowing for the accurate discovery of an additional cortical layer cluster in place of the artifact (**Figure 5D**).

Unlike dryspots, hangnail artifacts are more complex. Upon visual inspection of the mitochondrial ratio of hangnail artifacts (**Figure 4J**), we noticed that hangnails display low variance across spots compared to non-artifact domains. Taking advantage of this, we developed a QC method that successfully distinguishes hangnail artifacts based on the multiscale variance of mitochondrial ratio (**Figure 5E**). Hangnail artifacts are distinguishable in the first principal component of the multiscale variance (**Figures 5F-G**), and we found that *k*-means is superior to both Gaussian mixture models and density-based clustering non-parametric algorithms (DBSCAN) in clustering artifact from non-artifact spots (**Figure S5**). We additionally show that multiscale variance is superior to using a single neighborhood size for region-level artifact detection (**Figure S5**). Removing hangnail artifacts identified by SpotSweeper leads to improved downstream analyses, as evidenced by improved ranking of known L6 marker genes in one-vs-all spatial domain DE analyses (**Figure 5H**). These results highlight the importance of accurate artifact identification and removal in enhancing the reliability of SRT data.

## 3 Discussion

Quality control is vital across next-generation sequencing technologies to ensure data accuracy and integrity [7]. We present here the first spatially-aware QC methods developed specifically for SRT data. SpotSweeper improves spot-level quality control by using local *k*-nearest neighbor approaches to detect outliers within their spatial context, resulting in increased retention in high-quality spots. Using SpotSweeper, we discovered a set of spots in the 10x Genomics Visium platform with systematically low library size. We also characterized region-level artifacts unique to SRTs, and developed spatially-aware methods to detect and remove these artifacts. SpotSweeper can be used with spot-based SRTs to detect and remove both low quality spots and region-level artifacts prior to downstream analyses.

We demonstrated here that local outlier detection with SpotSweeper is superior to global outlier approaches commonly used for SRT data. Previous work has attempted to account for biological heterogeneity within snRNA-seq datasets by clustering nuclei based on their gene expression profiles prior to performing cell-level QC [28]. This can be computationally expensive for large datasets and is ineffective when low quality nuclei form a distinct cluster. We circumvent these issues by leveraging the spatial coordinates inherent to SRTs to normalize each spot based on their surrounding neighbors using a fast *k*-nearest neighbors algorithm. This ultimately increases the retention of high-quality spots, and thus, the statistical power for downstream analyses. We additionally characterized two distinct regional artifacts, dryspots and hangnails that are unique to SRT, and demonstrate that SpotSweeper is capable of accurately identifying these artifacts. This further improves clustering and marker gene detection, and is likely to have important implications for between-condition differential expression analyses (i.e., case vs control) [7, 29].

The proposed methods have some limitations that are open directions for future work. SpotSweeper is currently only compatible with spot-based SRT platforms, such as Visium. Image-based methods, such as MERFISH [30] and Xenium [31], profile the location of hybridized mRNA molecules with subcellular resolution. Regional artifacts due to tissue damage are also likely to remain a problem for image-based technologies. However, these technologies are limited to a smaller number of genes and thus typically do not include mitochondrial genes. While SpotSweeper currently uses multiscale variance of mitochondrial ratio to detect these artifacts, it is possible that a similar approach utilizing the negative control genes normally included in image-based methods may be useful for detecting damaged tissue sections. In addition, rasterization techniques that aggregate mRNA counts into spatial pixels [32] will increase compatibility with the current SpotSweeper workflow, while ensuring the scalability of our approaches for imaging-based platforms. Moreover, the current implementation of SpotSweeper should be amenable to future spot-based technological advancements, such as the VisiumHD platform from 10x Genomics [33], that have substantially increase spatial resolution with complete tissue coverage.

In summary, we propose the first spatially-aware QC methods that detect both spot- and regional-level artifacts for SRT data. These methods reduce bias due to biological heterogeneity and accurately identify regional artifacts that arise due to common tissue processing errors, improving both marker gene detection and spatial clustering steps. Our method is freely available at https://bioconductor.org/packa ges/SpotSweeper.

## 4 Methods

### 4.1 Preprocessing

The *SpotSweeper* workflow begins by using standard preprocessing steps. For the analyses performed here, we added spot-level QC metrics (i.e. library size, unique genes, and mitochondrial ratio) by using the addPerCellQCmetrics function from the *scuttle* R/BioConductor package. No gene expression data is otherwise used for in the *SpotSweeper* workflows.

### 4.2 Global QC methods

We used common QC workflows developed for snRNA-seq were implemented in the *scater* R/Bioconductor package. Individual spots were determined to be global outliers based on fixed thresholds (*<* 500 total transcripts, *<* 500 unique genes, or *>* 0.25 mitochondrial ratio) using the isOutlier function. The data were then summarized to the show the percent of discarded spots per manually annotated spatial domain for each sample, and visualized using the and *escheR* R/Bioconductor packages[34].

### 4.3 Spot-level algorithm and parameters

In contrast to global outliers, we define local outliers as spots that are outliers within their local neighbors in one or more of the three standard QC metrics (i.e., library size, unique genes detected, or mitochondrial ratio). Local neighborhoods are defined as the *k*-nearest neighbors[35] for each spot using their spatial co-ordinates using the BiocNeighbors R/Bioconductor package[36]. For Visium samples, we find that a neighborhood of three concentric circles (*k* =18) around each spot works well. Robust z-score transformation[17] of each spot is then used to normalize QC features across local neighborhoods. For each spot *i*, the robust z-score transformation can be formally defined as:

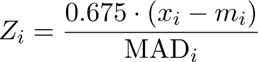

where *m_i_*is the median of the neighbors’ values:

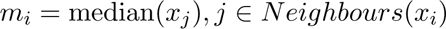

and the median absolute deviation (MAD) is defined as:

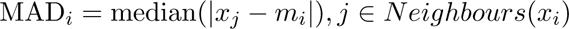

Adding the scaling factor of 0.675 makes the MAD comparable to the standard deviation under the assumption of normally distributed data. This in turn makes the robust z-score comparable to a standard z-score[18].

### 4.4 Detection and analysis of Visium spots with systematically low total UMI

Visium spots with systematically low total UMI (*n*=6 in total) were defined as spots that were detected as local outliers (*<* 3 z-scores) in over half (*>* n=22) of the total Visium samples used across Maynard et al. (n=12), Huuki-Myers et al. (n=30), and mouse coronal brain section (10x Genomics; n=1) datasets. Local outliers were detected using the localOutliers function from the *SpotSweeper* R/BioConductor package. The barcodes identifying these spots were then saved for further analyses.

### 4.5 Barcode sequence alignment and differential K-mer analysis

To better determine how barcode sequences may bias total UMI counts, we calculated the mean library size for each spot/barcode across all Visium samples (*n*=43) and the six barcodes with the highest mean UMI total were found. To compare the sequences of top mean-ranked barcodes and barcode with systemaic biased towards small library size, DNAStringSet objects were made using the barcodes with the *Biostrings* R/BioConductor package. We then aligned the sequences using the *msa*function from the *msa* R/BioConductor package.

All K-mer possibilities (e.g. A, AC, TGT, etc) for *k*= 1-4 were counted using the *Biostrings* R package. Differential *K*-mer testing between top and bottom mean-ranked barcode groups was carried out using student’s *t*-test. Volcano plots of differentially expressed *K*-mers were generated using the *EnhancedVolcano* R/CRAN package. Differential *K*-mers were further visualized using a heatmap generated by the pheatmap R package.

### 4.6 Artifact-level model and parameters

To find regional artifacts in the Huuki-Myers et al. dataset, the standard QC metrics (library size, unique genes, and mitochondrial ratio) for all samples were first visualized by generating spot plots using the *escheR* package. Visualization of hangnail artifacts revealed low variance in mitochondrial ratio (**Figure 4J**). Taking advantage of this, we developed a method to classify hangnail artifacts based on the multiscale variance. First, the local variance of mitochondrial ratio is computed at various scales (i.e., neighborhood sizes). To do this, multiple neighborhood sizes were defined as one to five concentric circles around each spot. For Visium, the exact neighborhood size, *K*, per concentric circle, *c*, can be defined as:

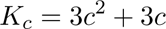

To assess the variability of mitochondrial content, we calculated the variance of the mitochondrial ratio within each defined local neighborhood. Preliminary analyses revealed that the mean mitochondrial ratio was a significant predictor of variance, suggesting a bias in local variance estimations related to the mean. To correct for this mean bias, we implemented robust linear regression using the iterative reweighted least squares algorithm[37]. This approach models the mean-variance relationship while being robust to the influence of outliers. We used the *rlm* function from the MASS package in R[38], applying default parameters. The residuals from this model provided an estimate of the local variance, adjusted for the mean mitochondrial ratio.

Following the estimation of mean-corrected local variance, principal component analysis is applied to the log-normalized local variances to reduce the dimensionality of the dataset; ultimately aiming to separate normal biological variance and variance induced by technical artifacts. We find that hangnail artifacts distinctly cluster in the first two principal components (**Figure 5F**). To classify these artifacts, we employed *k*-means clustering[39] with *k* =2. The classification of neighborhoods into artifact or non-artifact categories was subsequently determined by identifying the cluster with the lower average local variance. This cluster is automatically annotated as the artifact group. This is implemented in the findArtifacts function in the *SpotSweeper* package.

For the detection of dryspot artifact, the DBSCAN algorithm [40] was applied to the log2-transformed number of the UMI counts (library size) and log2-transformed number of unique genes. This was implemented using the dbscan function from the DBSCAN R/CRAN package[41] with the radius of the epsilon neighbor set to 0.5 (eps = 0.5) and the minimum number of points set to 20 (minPts = 20). Default parameters were otherwise used. We have implemented this procedure in the findDryspot function within the *SpotSweeper* package.

### 4.7 Spatially-aware clustering of spatial domains

Clustering of spatial domains (i.e., cortical layers) was achieved using the spatially-aware clustering method, BayesSpace[42]. To prepare the data, the mRNA counts for all samples were log-normalized using the logNormCounts function from the *scuttle*package. The mean-variance relationship was modeled using the modelGeneVar function prior to finding the the top 3000 highly-variable genes using the getTopHVGs function (both from the *scran* package), and dimensional reduction was performed with the top 3000 highly-variable genes using the runPCA function from the *scater* package. Spatially-aware clustering was then performed on the top 50 principal components using the spatialCluster function from the *BayesSpace* R/Bioconductor with 7 clusters (q = 7) and 10,000 iterations (nrep = 10000). This was conducted before and after artifact removal to determine the impact of discarding regional artifacts.

### 4.8 Differential expression analysis between spatial domains

To determine the rank of canonical marker genes before and after artifact removal in **Figure 5**, one vs all differential expression analyses were conducted for BayesSpace cluster #5 (shown in **Figure 4**; yellow) both before and after artifact removal using the findMarkers function from *scran*. The rankings of canonical L6 marker genes (*CCK*, *SCN3B*, *KRT17*, *SEMA3E*, *NR4A2*, *NTNG2*, and *SYNPR*)[10, 43, 44] were then compared.

### 4.9 Computational Implementation

*SpotSweeper* is implemented as an R package within the Bioconductor framework, using the *BiocNeighbors* package for local neighborhood detection, *stats* package for mean and variance calculations, *spatialEco* package for robust z-score normalization, *scater* for the implementation of principal component analysis, and *escheR* package for visualization. *SpotSweeper* takes advantage of the existing SpatialExperiment infrastructure for loading SRT input data and storing results. This allows for seemless integration in the existing Biocondcutor-based workflows.

### 4.10 Visium human DLPFC datasets

The (*n* = 12) Visium human DLPFC dataset from Maynard et al. consists of twelve total samples from three different neurotypical donors, measured with the 10x Genomics Visium platform[10]. The dataset was originally published by Maynard et al. and subsequently released through the spatialLIBD R/Bioconductor package. The data used in this manuscript were acquired using the fetch data function with type set to “spe”. This dataset contains transcriptome-wide gene expression measurments across 47,681 spots under tissue areas. The data were manually annotated with labels for the six cortical layers and white matter in the original study, which we use as an approximate for ground truth labels for method evaluation. These data do not contain regional artifacts such as hangnails or dryspots.

The (*n* = 30) Visium human DLPFC dataset from Huuki-Myers et al. consists of thirty total samples from ten different neurotypical donors, measured with the 10x Genomics Visium platform and made published by Huuki-Myers et al[9]. The processed data is also available via the fetch data function from the *spatialLIBD* package. This datasets contains transcriptome-wide gene expression measurements across 118,800 spots under with tissue areas. In this manuscript, we are especially interested in the dryspot and hangnail artifacts present in samples “Br3942 mid” and “Br8325 ant”, respectively.

### 4.11 Mouse Coronal Brain dataset

The mouse brain dataset consists of a single coronal section measured with the Visium platform, generated by 10x Genomics. This dataset is publicly available from 10x Genomics. For the analyses in this manuscript, this was acquired via the *STexampleData* R/Bioconductor package using the *Visium mouseCoronal* function. This dataset also contains transcriptome-wide gene expression data across 2,702 spots under tissue areas.

## Data availability

The DLPFC datasets used for analyses in this manuscript can be obtained from spatialLIBD (http://research.libd.org/spatialLIBD) in SpatialExperiment format, which includes Manual Annotation labels from the original sources. All other data supporting the findings of this study are available within the article and its supplementary files. Any additional requests for information can be directed to, and will be fulfilled by, the lead contact.

## Code availability

The code that generates these figures is deposited at https://github.com/boyiguo1/Manuscript-SpotSweep er (Zenodo DOI: 10.5281/zenodo.11489067). The open source software package SpotSweeper is available in the R programming language and freely available on Bioconductor (https://bioconductor.org/package s/SpotSweeper). We used SpotSweeper version 0.99.5 for the analyses in this manuscript.

## Abbreviations

DLPFC: dorsolateral prefrontal cortex
MAD: median absolute deviation
QC: quality control
sc/snRNA-seq: single-cell/nucleus RNA-sequencing
SRT: spatially-resolved transcriptomics
WM: white matter

## Author contributions

- **M.T.**: Conceptualization, Methodology, Software, Validation, Formal analysis, Investigation, Data curation, Writing, Visualization
- **S.C.H.**: Conceptualization, Resources, Writng - Review & Editing, Visualization, Supervision, Project administration, Funding Acquisition
- **B.G.**: Conceptualization, Writng - Review & Editing, Visualization, Supervision, Project administration

## Acknowledgements

We would like to thank our collaborators at the Lieber Institute for Brain Development, especially Dr. Keri Martinowich, for valuable input and feedback during the development and application of these methods to identify the nature and cause of regional artifacts during sample preparation. We would like to also thank the maintainers of the Joint High Performance Computing Exchange (JHPCE) compute clusters at Johns Hopkins Bloomberg School of Public Health for providing essential computing resources.

## Supplementary Materials

### Supplemental Figures S1-S5

**Supplementary Figure S1:**
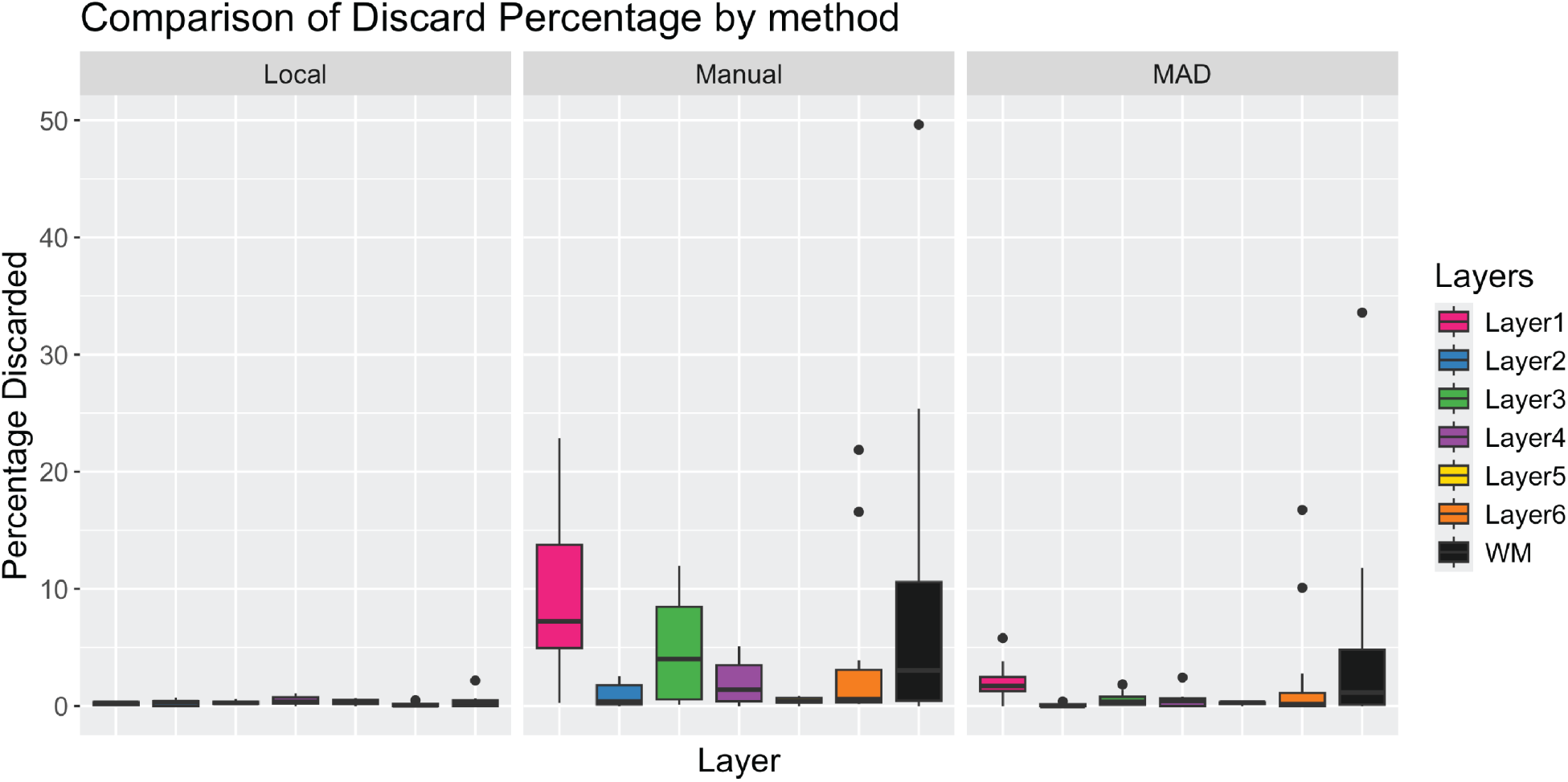
Comparison of commonly used spot-level QC methods for SRT data using the *n*=12 Maynard et al. [10] Visium samples. Three different QC approaches were considered: local outliers (SpotSweeper) (left), global outliers using fixed thresholding (middle), and global outliers using a threshold of three MADs based on the sample-wise distributions of outliers of each mitochondrial ratio, library size, and unique genes (right). Figure is showing boxplots of the percentage of discarded spots per tissue sample (a point in the boxplot) stratified by the cortical layers: white matter and layers 1-6.

**Supplementary Figure S2:**
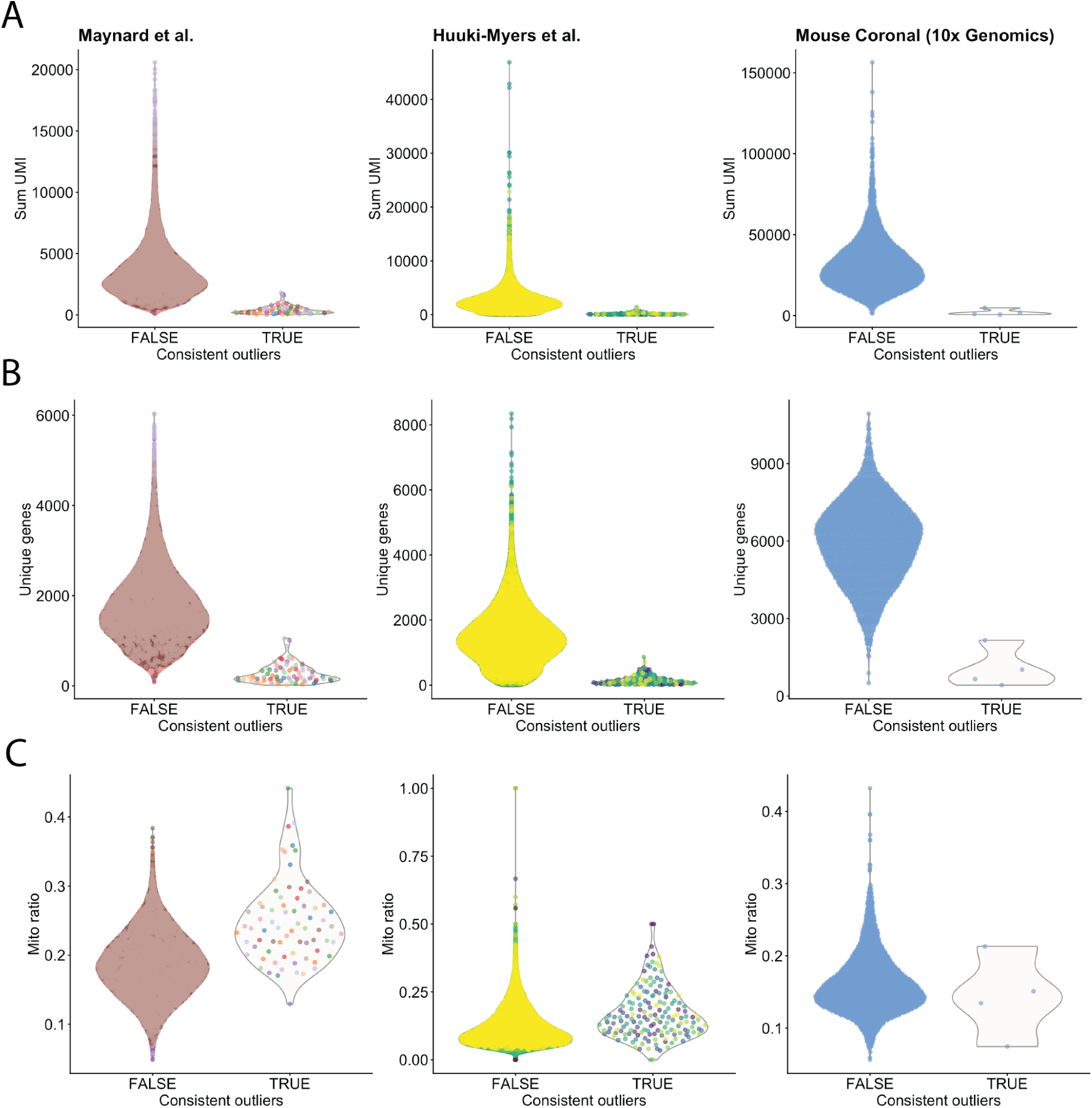
Violin plots showing standard QC metrics library size (A), unique genes (B), and mitochondrial ratio (C) for the six spots that are consistent outliers across Maynard et al. (*n*=12), Huuki-Myers et al. (*n*=30), and mouse coronal section datasets (*n*=1). All plots are color by sample. Only spots underlying tissue samples were included.

**Supplementary Figure S3:**
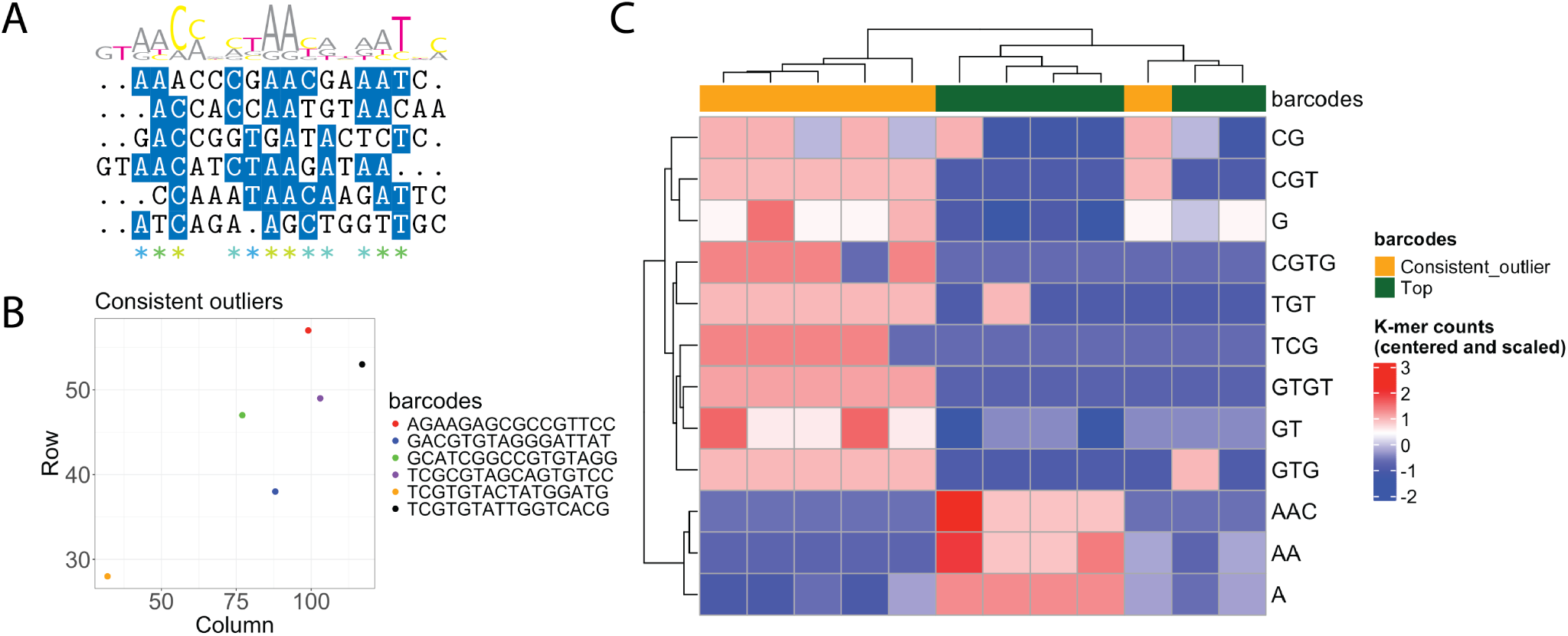
(A) Sequence alignment of the top six mean-ranked barcodes reveals relatively little homology across barcodes. (B) The barcode sequences and spatial coordinates of the six consistent Visium outlier spots. (C) Heatmap of centered and scaled counts for differentially expressed (*p* ¡ .05) *k*-mers between consistent outliers and the top six mean-ranked barcodes. Hierarchical clustering of the columns and rows nearly perfectly distinguished outliers from top ranked barcodes, demonstrating substantial homology within the groups.

**Supplementary Figure S4:**
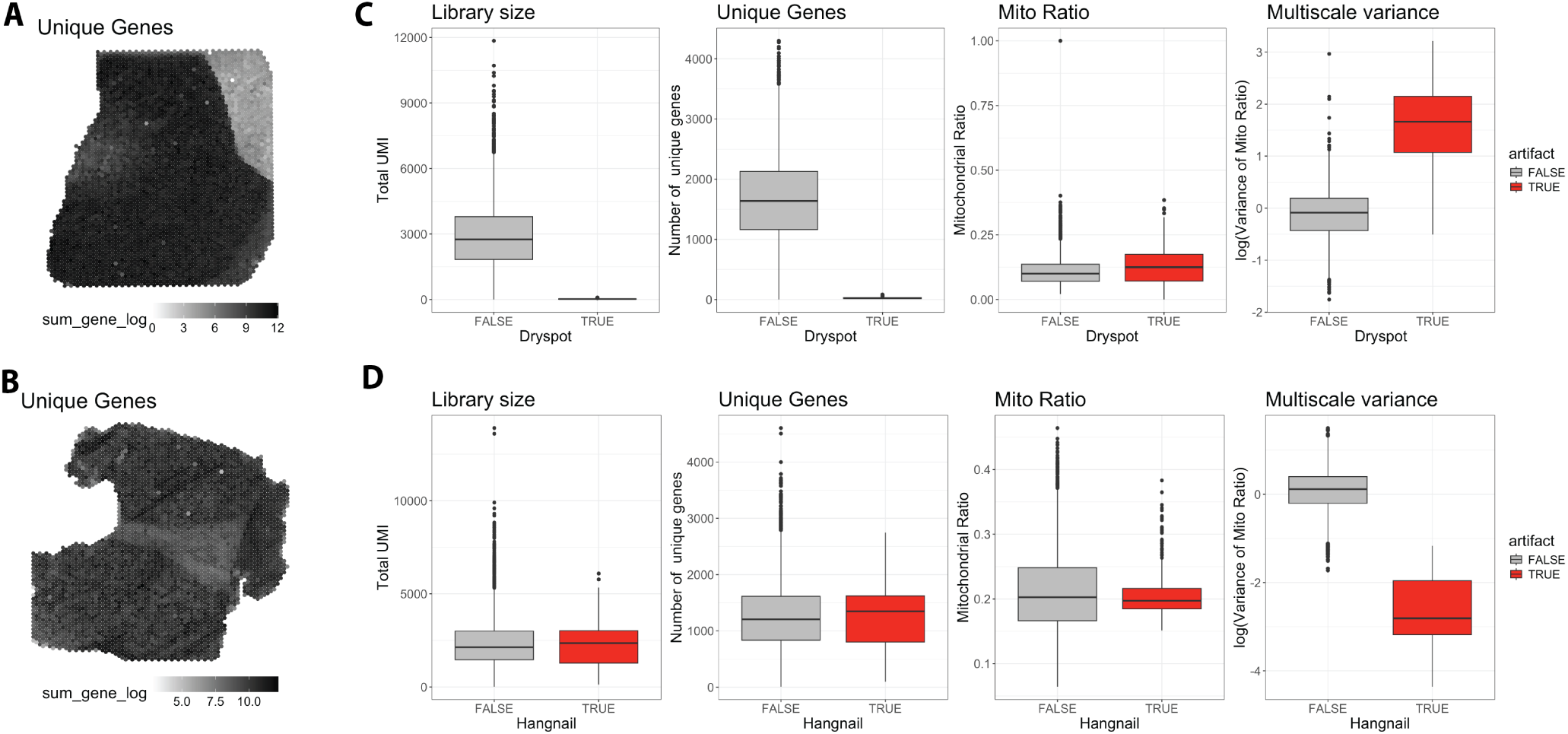
Spot plots showing that the number of unique genes (log-transformed) in dryspot (A) and hangnail artifacts (B). (C) Box plots demonstrating that dryspot artifacts display lower library size and uniques genes, but no difference in average mitochondrial ratio. Dryspots do display higher average multiscale variance in mitochondiral ratio. (D) Box plots demonstrating that hangnail artifacts display no mean differences in library size, number of unique genes, or mitochondrial ratio. Hangnails do, however, display lower average multiscale variance in the mitochondrial ratio.

**Supplementary Figure S5:**
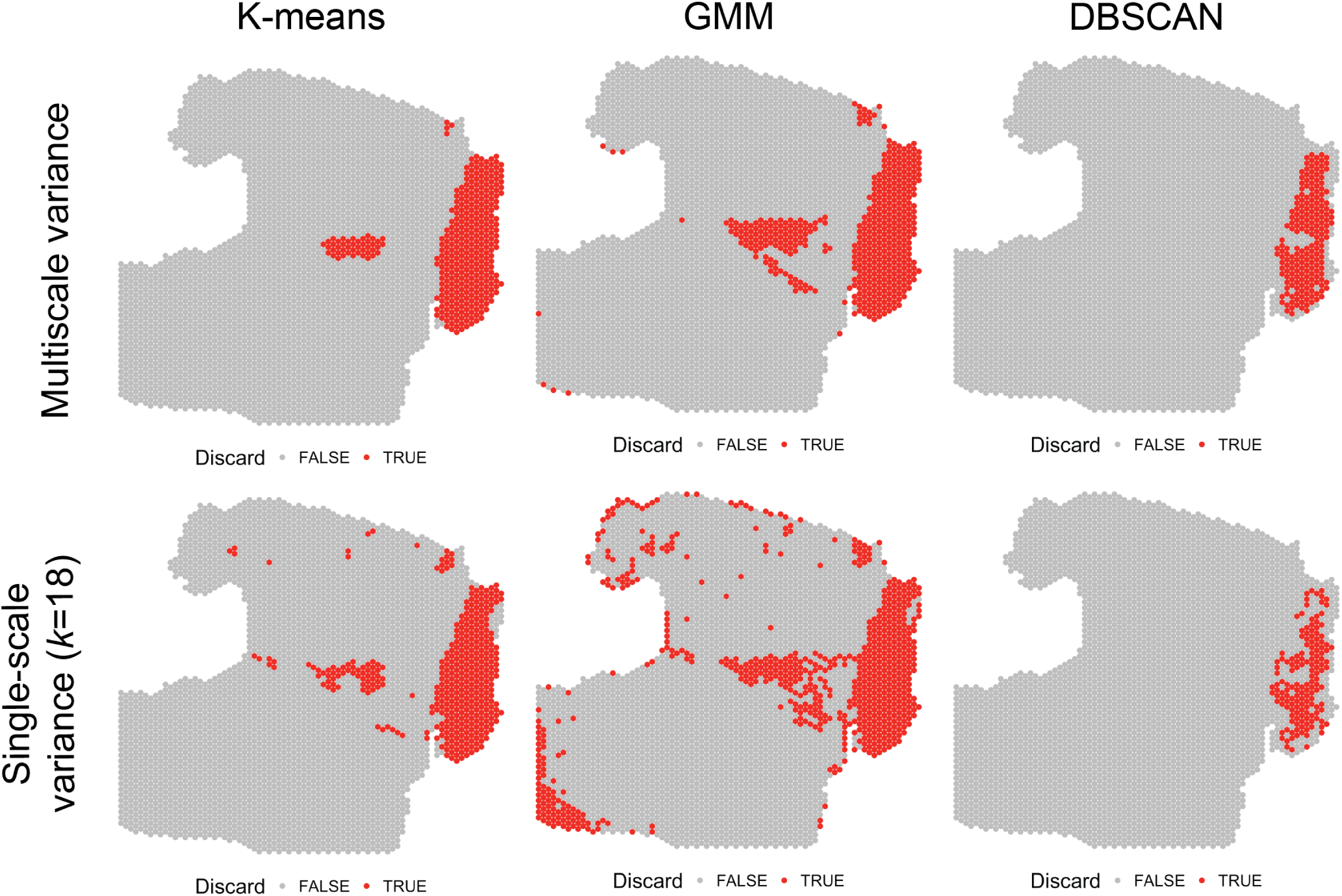
(Rows) Comparison of hangnail detection using single-scale (*k* = 18; one concentric circle per spot) versus multiscale variance (1-5 concentric cicles per spot). (Columns) Comparison of (*k*-means (*k* = 2), Gaussian mixed models (*k* = 2), and DBSCAN methods for accurately clustering hangnail artifacts.

